# Efficient and cost-effective bacterial mRNA sequencing from low input samples through ribosomal RNA depletion

**DOI:** 10.1101/2020.06.19.162412

**Authors:** Chatarin Wangsanuwat, Kellie A. Heom, Estella Liu, Michelle A. O’Malley, Siddharth S. Dey

## Abstract

RNA sequencing is a powerful approach to quantify the genome-wide distribution of mRNA molecules in a population to gain deeper understanding of cellular functions and phenotypes. However, unlike eukaryotic cells, mRNA sequencing of bacterial samples is more challenging due to the absence of a poly-A tail that typically enables efficient capture and enrichment of mRNA from the abundant rRNA molecules in a cell. Moreover, bacterial cells frequently contain 100-fold lower quantities of RNA compared to mammalian cells, which further complicates mRNA sequencing from non-cultivable and non-model bacterial species. To overcome these limitations, we report EMBR-seq (Enrichment of mRNA by Blocked rRNA), a method that efficiently depletes 5S, 16S and 23S rRNA using blocking primers to prevent their amplification, resulting in greater than 80% of the sequenced RNA molecules from an *E. coli* culture deriving from mRNA. We demonstrate that this increased efficiency provides a deeper view of the transcriptome without introducing technical amplification-induced biases. Moreover, compared to recent methods that employ a large array of oligonucleotides to deplete rRNA, EMBR-seq uses a single oligonucleotide per rRNA, thereby making this new technology significantly more cost-effective, especially when applied to varied bacterial species. Finally, compared to existing commercial kits for bacterial rRNA depletion, we show that EMBR-seq can be used to successfully quantify the transcriptome from more than 500-fold lower starting total RNA. Thus, EMBR-seq provides an efficient and cost-effective approach to quantify global gene expression profiles from low input bacterial samples.

## Introduction

Bacterial species pervade our biosphere and millions of years of evolution have optimized these microbes to perform specific biochemical reactions and functions; processes that could potentially be adapted to develop a variety of products, such as renewable biofuels, antibiotics, and other value-added chemicals [1–5]. Bacterial messenger RNA (mRNA) sequencing provides a snapshot of the genome-wide state of a microbial population, and therefore enables fundamental understanding of these varied microbial functions and phenotypes [6].

However, compared to eukaryotes, mRNA sequencing from bacterial samples has been more challenging for several reasons. First, unlike in eukaryotes, bacterial mRNA does not contain a poly-A tail at the 3’ end that can be used to easily enrich for these molecules during reverse transcription [7, 8]. Further, total RNA isolated from bacterial cells typically contains greater than 95% ribosomal RNA (rRNA), and therefore cost-effective and high coverage sequencing of the transcriptome requires the development of efficient strategies to deplete the abundant 5S, 16S and 23S rRNA molecules [9]. Finally, bacterial cells typically contain approximately 100-fold lower RNA than mammalian cells, and as the starting amount of total RNA when working with rare, non-cultivable, and non-model bacterial species can be limiting, it is a challenge to robustly and accurately capture the transcriptome from small quantities of total RNA with minimal amplification biases [10].

Several commercial kits have been developed to deplete bacterial rRNA from total RNA samples, including the MICROBExpress Bacterial mRNA Enrichment Kit (Thermo Fisher Scientific), the RiboMinus Transcriptome Isolation Kit, bacteria (Thermo Fisher Scientific), and the Ribo-Zero rRNA Depletion Kit (Illumina) [11]. These techniques rely on subtractive hybridization to deplete rRNA and typically work at a scale of hundreds of nanograms to micrograms of starting total RNA. Further, as these commercial kits are only effective on species targeted in the standard probe set, it is challenging to extrapolate these methods to diverse bacterial species [9, 11]. While this limitation of pre-designed kits have been overcome through the development of workflows to generate custom subtractive hybridization probe sets for any species of interest, they still operate at microgram quantities of starting material and either require multiple rounds of hybridization or a series of oligo optimization steps prior to optimal performance [12, 13]. An alternate approach relies on the Terminator™ 5’-phosphate-dependent exonuclease (TEX) (Lucigen) to specifically degrade rRNAs with 5’-monophosphate ends but not mRNAs with 5’-triphosphate ends; however, this method typically has lower efficiencies than other existing rRNA depletion strategies [10, 14, 15]. A more recent method uses complementary single-stranded DNA probes to tile rRNAs that are subsequently degraded by RNase H [16]. The commercial NEBNext Bacteria rRNA depletion kit (NEB) employs a similar strategy and can be applied to as low as 10 ng of starting total RNA. Similarly, another approach uses a pool of tiled single-guide RNAs to direct Cas9 mediated cleavage of rRNA-derived cDNA to deplete rRNA while another approach uses targeted reverse transcription primers designed to avoid capturing rRNAs [17, 18]. However, all these methods require a large array of probes that can be expensive to synthesize and potentially need to be redesigned for distant bacterial species [16–18].

Therefore, in this work we have developed EMBR-seq (Enrichment of mRNA by Blocked rRNA), a new technology that overcomes the limitations of sequencing mRNA from bacterial samples by: (1) Using 5S, 16S and 23S rRNA blocking primers and poly-A tailing to specifically deplete rRNA and enrich mRNA during downstream amplification; (2) Using a single blocking primer for each of the three abundant rRNA molecules, thereby enabling rapid adaptation to different bacterial species and significantly reducing the cost per sample; and (3) Using a linear amplification strategy to amplify mRNA from as low as 20 picograms of total RNA with minimal amplification biases. We applied EMBR-seq to a model *E. coli* system to demonstrate efficient mRNA enrichment and sequencing with increased sensitivity in gene detection. Further, we show that our method accurately captures the genome-wide gene expression profiles with minimal technical biases. Thus, EMBR-seq is an efficient and cost-effective approach to sequence mRNA from low-input bacterial samples.

## Results

### EMBR-seq uses blocking primers to deplete rRNA

To overcome the limitations described above, we developed EMBR-seq, a new technique to efficiently deplete rRNA from total RNA, thereby enabling cost-effective sequencing of mRNA from bacterial cells. To minimize rRNA-derived molecules in the final sequencing library, we first incubated the total RNA with rRNA blocking primers, designed specifically to bind the 3’ end of 5S, 16S and 23S rRNA, followed by poly-adenylation with *E. coli* poly-A polymerase (Fig. 1 and Methods). To deplete rRNA, EMBR-seq only requires primers at the 3’ end of rRNA, unlike recent methods that tile oligonucleotides along the entire length of rRNA molecules, thereby significantly reducing costs and making our approach more easily translatable to other bacterial species. The blocking primers generate double-stranded RNA-DNA hybrid molecules at the 3’ end of rRNAs, which reduces subsequent poly-adenylation and downstream amplification of rRNA molecules, as the poly-A polymerase preferentially adds adenines to single-stranded RNA [19]. Thereafter, the reaction mixture is reverse transcribed following the addition of a poly-T primer. This primer has an overhang containing a sample-specific barcode to enable rapid multiplexing and reduction in library preparation costs, the 5’ Illumina adapter, and a T7 promoter [20]. After second strand synthesis, cDNA molecules are amplified by *in vitro* transcription (IVT). However, as only cDNA molecules deriving from a poly-adenylated RNA have a T7 promoter, our technique further amplifies mRNA-derived molecules for sequencing whereas rRNA-derived molecules are excluded from IVT amplification. The amplified RNA from IVT is then used to prepare Illumina sequencing libraries, as described previously (Fig. 1 and Methods) [20–22].

**Figure 1:**
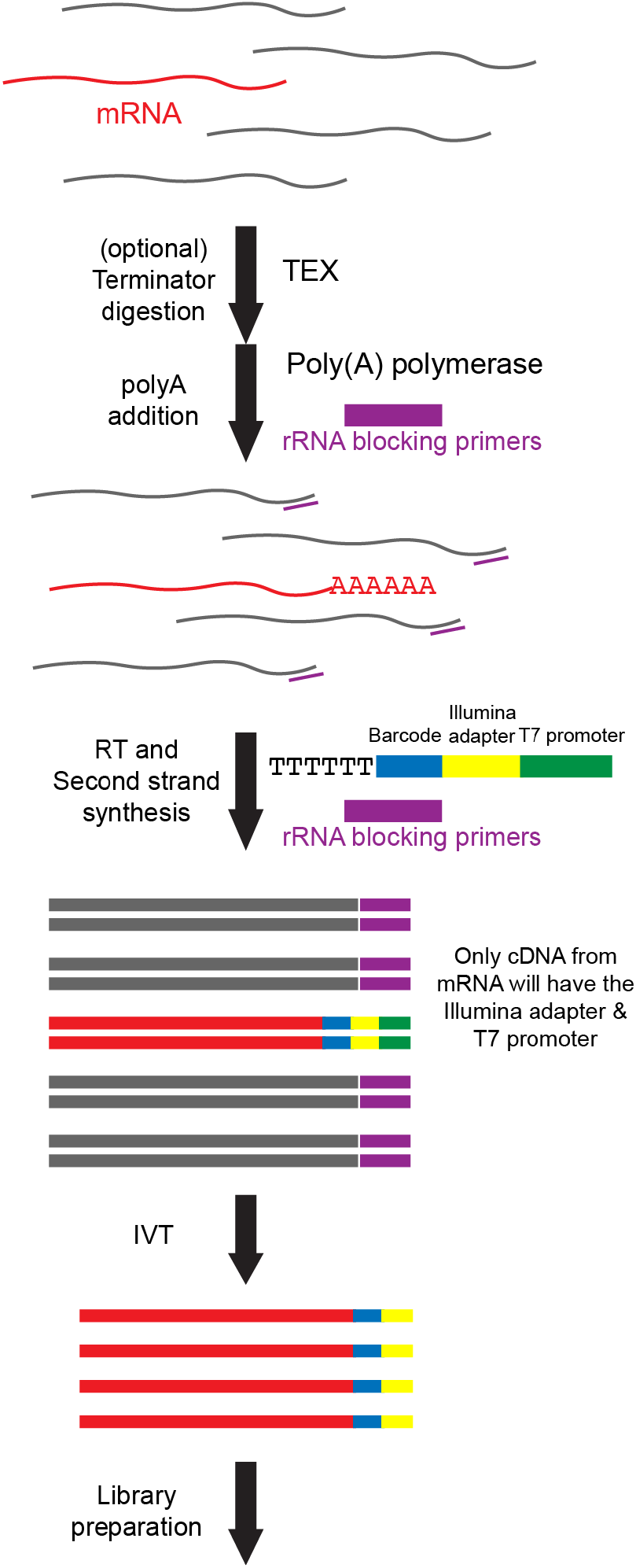
Schematic of EMBR-Seq. After performing an optional Terminator™ 5’-phosphate-dependent exonuclease digestion, poly(A) polymerase and rRNA blocking primers (purple) are added to total bacterial RNA (mRNA in red and rRNA in gray). Blocking primers specifically bind to the 3’ end of 5S, 16S, and 23S rRNAs, resulting in the preferential addition of a poly-A tail to mRNA molecules. Next, reverse transcription is performed using (i) a poly-T primer, which has an overhang containing a sample-specific barcode (blue), 5’ Illumina adapter (yellow), and T7 promoter (green), and (ii) rRNA blocking primers to convert poly-adenylated RNA and rRNA molecules, respectively, to cDNA. The cDNA molecules are then amplified by *in vitro* transcription, and the amplified RNA is used to prepare Illumina libraries. As the rRNA-derived cDNA does not contain a T7 promoter, these molecules are not amplified during *in vitro* transcription, resulting in rRNA depletion.

### EMBR-seq efficiently depletes rRNA to sequence bacterial mRNA

We applied EMBR-seq to total RNA isolated from the exponential growth phase of *E. coli* strain K12 (MG1655). Starting from 100 ng of total RNA, we were able to successfully make Illumina libraries that were sequenced and mapped to the *E. coli* transcriptome. While total RNA from *E. coli* has previously been reported to consist of 95% rRNA [9], our control samples with no blocking primers had approximately 64% rRNA, consistent with previous observations that mRNA molecules are preferentially poly-adenylated compared to rRNA even in the absence of any blocking primers (Fig. 2a) [23]. Importantly, compared to the control samples, we observed a significant increase in rRNA depletion efficiency (from 64% to 16%), with 84% of the mapped reads corresponding to mRNA in samples treated with blocking primers (Fig. 2a). These results demonstrate that EMBR-seq achieves a level of mRNA enrichment that is better or comparable to recent bacterial rRNA depletion reports [11–13, 15–18].

**Figure 2:**
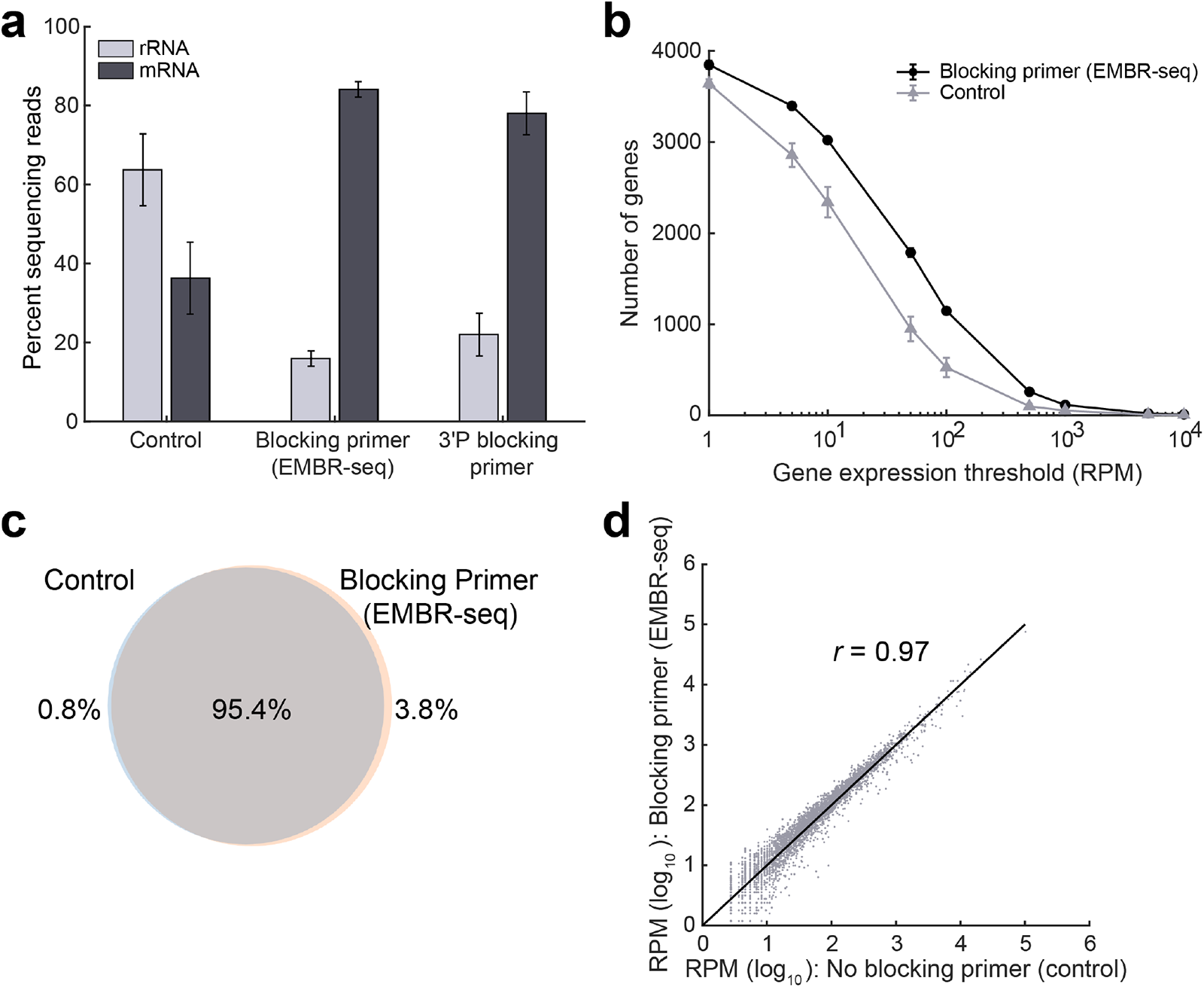
Blocking primers in EMBR-seq deplete rRNA and provide a deeper view of the transcriptome without introducing technical biases. (**a**) In the presence of blocking primers, a 4-fold rRNA depletion and more than 2-fold mRNA enrichment is achieved compared to control samples. With the introduction of blocking primers in EMBR-seq, mRNAs account for more than 80% of the mapped reads, which is a greater than 16-fold increase compared to total RNA in *E. coli* cells. The 3’ phosphorylated blocking primers display similar but slightly lesser mRNA enrichment (*n* ≥ 2 replicates for all conditions). (**b**) Comparison between EMBR-seq and control samples in the number of genes detected above different expression thresholds (n = 3 for both conditions). For the EMBR-seq group, error bars are of the same scale as the size of the data points (**c**) Venn diagram shows that more than 99% of the genes detected in the control samples were also detected when using blocking primers in EMBR-seq. 99.2 % of all detected genes were found in the EMBR-seq samples and 96.2% in the control samples. The number of genes detected were calculated by combining data obtained from three control samples and three EMBR-seq processed samples. (**d**) Gene transcript counts with and without blocking primers are highly correlated (Pearson *r* = 0.97) suggesting that EMBR-seq does not introduce technical artifacts in quantifying gene expression (n = 3 for both datasets). These experiments were performed starting with 100 ng total RNA from *E. coli*. Error bars in panels (a) and (b) represent standard deviations.

We also tested modified blocking primers with a 3’ phosphorylation, designed to prevent Superscript II from reverse transcribing rRNA molecules. As expected, we observed rRNA depletion in these samples as well (from 64% to 22%), with 78% of the mapped reads corresponding to mRNA (Fig. 2a). However, compared to the unmodified blocking primers, these phosphorylated blocking primers were slightly less efficient at rRNA depletion (Fig. 2a). As the 3’ phosphorylated primers prevent polymerase extension, we hypothesize that the reduced rRNA depletion efficiency arises from the small fraction of rRNA molecules that get poly-adenylated, primed by the poly-T primers, and copied through the short 30 bp RNA-DNA hybrid due to the strand-displacement activity of the reverse transcriptase. Therefore, given the reduced efficiency and higher costs of the 3’ phosphorylated blocking primers, all further experiments were performed with unmodified blocking primers.

As an alternate strategy, we also incorporated TEX treatment in EMBR-seq as it has previously been shown to specifically degrade rRNAs with 5’-monophosphate ends but not mRNAs that have 5’-triphosphate ends [10, 14, 15]. While we again observed rRNA depletion and a corresponding enrichment of mRNA compared to control samples, the effects were less pronounced with a less than 2-fold rRNA depletion, consistent with previous reports (Fig. S1) [14, 15]. We hypothesize that this reduced efficiency arises from RNA degradation that might occur during the incubation at 37°C for 1 hour or the additional cleanup step that is necessary prior to treatment with the poly-A polymerase. As a result, we find that blocking primers alone provide the most significant rRNA depletion and mRNA enrichment, and therefore all further experiments were performed without TEX treatment.

### EMBR-seq is a cost-effective bacterial mRNA sequencing technology

In designing the steps of EMBR-seq, we wanted to develop a method that is both easily applied and cost-effective. Due to its simplicity, the cost per rRNA depletion reaction in EMBR-seq is ~$0.40, which is at least an order of magnitude lower than other recent rRNA depletion methods and commercial kits [11–13, 15–18] (Fig. S2a and Supplementary Table 1). The total cost of EMBR-seq, starting from total bacterial RNA to the final Illumina library, was estimated to be ~$36 per sample. However, the total cost per sample decreases as more samples are multiplexed in the same Illumina library. For example, when 96 samples are multiplexed, the cost per sample drops to ~$20, primarily due to the pooling of samples after second-strand synthesis that then requires only a single IVT and Illumina library preparation reaction downstream (Fig. S2b). Thus, EMBR-seq is a simple and cost-effective approach to sequence mRNA from total bacterial RNA.

### EMBR-seq provides a detailed view of the transcriptome without introducing technical biases

Next, we systematically compared the gene expression profiles obtained from control and rRNA depleted samples to investigate if the use of blocking primers provides a deeper view of the transcriptome without introducing technical artifacts. First, after downsampling sequencing reads to the same depth, we detected 3628 genes in the control samples, while in the mRNA enriched samples we detected 3852 genes, with 99% of the genes in the control samples also detected in the mRNA enriched samples (Fig. 2b,c). Moreover, at different levels of downsampling, we detected more genes using EMBR-seq compared to the control samples (Fig. S3). This suggests that we can measure the genome-wide gene expression landscape in a more cost-effective way using EMBR-seq. Further, the number of genes detected above different expression thresholds was consistently higher for the mRNA enriched samples compared to the control samples (Fig. 2b). This shows that EMBR-seq is able to detect more genes at different gene expression levels, spanning over three orders of magnitude. Finally, we observed that gene expression between the control and mRNA enriched samples were highly correlated (Pearson *r* = 0.97) revealing that the blocking primers do not introduce technical biases in the quantification of gene expression (Fig. 2d). Collectively, these results demonstrate that our new cost-effective method is able to accurately capture the transcriptome of bacterial cells.

### EMBR-seq allows mRNA sequencing from low input total RNA

In many practical applications involving non-model and non-cultivable bacterial species, the starting amount of total RNA available for RNA sequencing can be limiting. Therefore, we evaluated if we can successfully deplete rRNA and quantify gene expression from lower amounts of input material. We applied EMBR-seq to 20, 2, 0.2 and 0.02 ng of starting total RNA isolated from the exponential growth phase of *E. coli* strain K12. These starting quantities of total RNA were chosen as they are typically below the sensitivity and detection limit of commercial kits and previously reported methods [11, 17]. As before, we observed a greater than 3-fold depletion of rRNA across the range of input starting material, including at the lowest starting amount of 0.02 ng total RNA, with greater than 77% of the reads in the sequencing library deriving from mRNA molecules (Fig. 3a). Similarly, we observed that the total number of genes detected is higher than that in the control samples and is unaffected by the starting input amount of total RNA, except at the lower starting amounts of 0.2 ng and 0.02 ng total RNA (Fig. 3b). Finally, we also observed that gene expression was highly correlated between different amounts of starting total RNA (Fig. 3c and Fig. S4). These experiments conclusively demonstrate that we can successfully apply EMRB-seq to quantify gene expression from total RNA starting as low as 20 pg.

**Figure 3:**
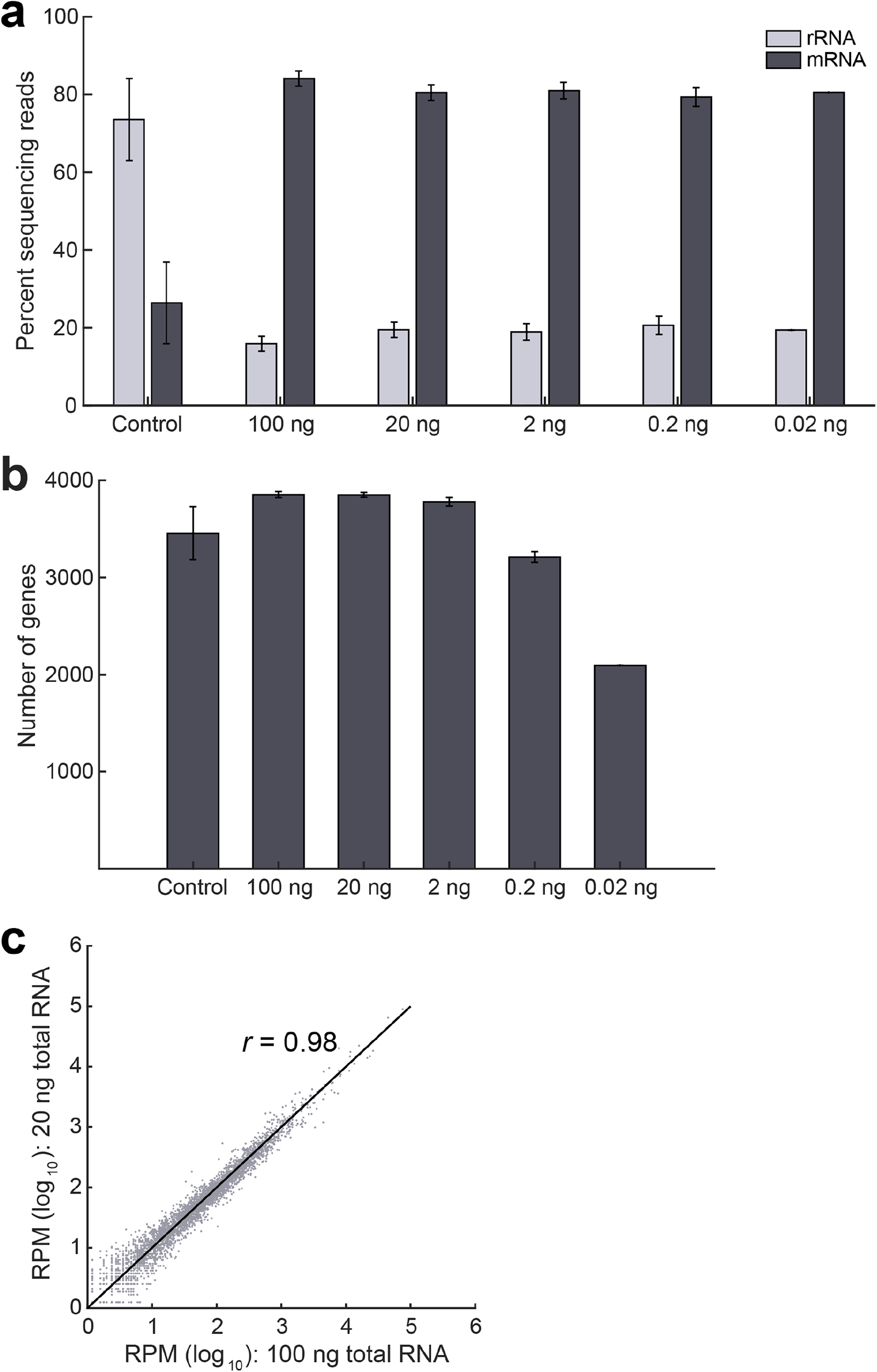
EMBR-seq can quantify the transcriptome from low input total RNA. (**a**) Similar levels of rRNA depletion and mRNA enrichment are observed when the starting amount of total RNA is decreased from 100 ng to 0.02 ng (*n* ≥ 2, except at 0.02 ng where *n* = 1). The control represents average data of control samples made from different input levels of total RNA. The 100 ng data is reproduced from Figure 2a. (**b**) Compared to the control samples, more genes are detected when starting with at least 2 ng input total RNA. Fewer genes are detected when starting total RNA decreases to 0.02 ng. (**c**) Gene transcript counts are highly correlated (Pearson *r* = 0.98) between 100 ng and 20 ng input total RNA in EMBR-seq. Datasets from lower starting total RNA are also well correlated to the 100 ng samples (Fig. S4).

## Discussion

We have developed a new technology, EMBR-seq, to efficiently deplete rRNA from total RNA, thereby enabling a deeper view of the genome-wide distribution of mRNA in bacterial samples. Sequencing bacterial mRNA poses several challenges; for example, the inability to easily enrich mRNA that typically makes up less than 5% of total RNA and the limiting starting amounts of total RNA that may be available when working with non-cultivable bacterial samples [7–9]. Through the use of a single blocking primer per rRNA species, EMBR-seq efficiently minimizes the downstream amplification of rRNA molecules, thereby enabling a 4-fold depletion of rRNA in the final sequencing library (Fig.1 and 2a). As demonstrated in this work, the design of blocking primers at the 3’ end of rRNA molecules efficiently depletes rRNA from high quality total RNA samples; however, certain practical applications can produce degraded and fragmented RNA in which rRNA molecules will be less effectively depleted. We hypothesize that a straightforward strategy to overcome this challenge in EMBR-seq in the future is to design additional blocking primers, potentially 3-5 primers per rRNA species, that span the transcript length to minimize amplification of degraded rRNA molecules.

Starting with total RNA from *E. coli*, we show that efficient depletion of rRNA by EMBR-seq provides higher coverage of the transcriptome at the same sequencing depth (Fig. 2b and Fig. S3). For example, compared to the control samples, the number of unique genes detected increases from 3628 to 3852 in EMBR-seq (Fig. 2b). In particular, EMBR-seq improves detection of lowly expressed genes below 500 RPM (Fig. 2b). Further, EMBR-seq provides a more in-depth view of the transcriptional landscape without introducing technical artifacts. We find that 99% of the genes detected in the control group are also detected by EMBR-seq, and that gene expression levels between the two groups are highly correlated (Fig. 2c,d).

As EMBR-seq uses a single blocking primer per rRNA species, it is likely easily adaptable to other microbial species. Recent approaches that employ a large array of probes also achieve a high efficiency of rRNA degradation; however, the need to generate such a large pool of molecules makes it more challenging to extrapolate these methods to evolutionarily distant bacterial species compared to EMBR-seq [16–18]. In addition, the use of just one primer per rRNA species combined with the high level of sample multiplexing reduces cost significantly compared to other methods, enabling cost-effective and high-throughput processing of hundreds of samples simultaneously (Fig. S2). Finally, beyond rRNA, the approach used in EMBR-seq can potentially also be used to target other high abundance transcripts in total RNA.

We also demonstrated that EMBR-seq enables mRNA sequencing of low input RNA samples below the detection limit of commercial kits (Fig. 3a). Bacterial populations frequently contain diverse species, and even isogenic systems have been shown to display substantial cell-to-cell heterogeneity in gene expression that can give rise to dramatic cellular phenotypes [24–29]. Therefore, scaling down bacterial mRNA sequencing techniques to a single-cell level will enable quantification of this variability and provide a better understanding of how transcriptomic heterogeneity regulates cellular function [30, 31]. Over the last few years, a limited number of approaches have been developed to sequence the transcriptome of single bacterial cells. Early proof-of-concept methods were low throughput techniques that sequenced less than 10 single cells and generally suffered from significant technical noise [10, 32, 33]. More recently, Blattman *et al*. employed combinatorial barcoding to circumvent single cell isolation, enabling high throughput single-cell sequencing of bacterial cells [34]. However, this method did not deplete rRNA, resulting in low mRNA detection efficiencies of ~0.5-2% (or ~40 mRNA per *E. coli* cell). In another study, Kuchina *et al*. combined rRNA depletion with combinatorial barcoding to achieve ~5-10% mRNA detection efficiencies in *B. subtilis* [14]. These initial efforts suggest that improved methods could significantly advance single-cell mRNA sequencing in bacteria. EMBR-seq can successfully sequence mRNA from as low as 20 pg of total RNA; therefore, we anticipate that by coupling our rRNA depletion strategy with recent combinatorial barcoding techniques, we will be able to extend EMBR-seq to a single-cell resolution in the future [14, 34].

## Methods

### Bacterial strains and culture conditions

*Escherichia coli* MG1655 (ATCC: 700926) overnight cultures were inoculated into fresh LB medium at 1:50 and grown at 37°C with shaking (150 rpm). Upon reaching the exponential growth phase, the culture was centrifuged at 3000 g for 10 min. The media was removed and the pellet was resuspended in PBS to a concentration of 10^7^ cells per μL. The cells were stored on ice and total RNA extraction was performed immediately.

### RNA extraction

Trizol (Thermo Fisher Scientific, Cat. # 15596018) RNA extraction was performed following the manufacturer’s protocol. Briefly, 10^8^ cells were added to 750 μL Trizol, mixed, and then combined with 150 μL chloroform. After centrifugation, the clear aqueous layer was recovered and precipitated with 375 μL of isopropanol and 0.67 μL of GlycoBlue (Thermo Fisher Scientific, Cat. # AM9515). The pellet was washed twice with 75% ethanol and after the final centrifugation, the resulting pellet was resuspended in RNase-free water.

### EMBR-seq

#### Poly adenylation

100 ng of total RNA in 2 μL was combined with 3 μL poly(A) mix, comprised of 1 μL 5x first strand buffer [250 mM Tris-HCl (pH 8.3), 375 mM KCl, 15 mM MgCl2, comes with Superscript II reverse transcriptase, Invitrogen Cat. # 18064-014], 1 μL blocking primer mix (50 μM) (see *Primers*), 0.8 μL nuclease-free water, 0.1 μL 10 mM ATP, and 0.1 μL *E. coli* poly(A) polymerase (New England Biolabs, Cat. # M0276S). The mixture was incubated at 37°C for 10 min. In the control group, no blocking primers were added and 1.8 μL of nuclease-free water was added instead. The blocking primer mix was prepared by mixing equal volumes of 50 μM blocking primers specific to 5S, 16S, and 23S rRNA.

#### Reverse transcription

The polyadenylation product was mixed with 0.5 μL 10 mM dNTPs (New England Biolabs, Cat. # N0447L), 1 μL reverse transcription primers (25 ng/μL, see *Primers*), and 1.3 μL blocking primer mix (50 μM), and heated to 65°C for 5 min, 58°C for 1 min, and then quenched on ice. In the control samples, the blocking primers were again replaced with nuclease-free water. Next, 3.2 μL RT mix, consisting of 1.2 μL 5x first strand buffer, 1 μL 0.1 M DTT, 0.5 μL RNaseOUT (Thermo Fisher Scientific, Cat. #10777019), and 0.5 μL Superscript II reverse transcriptase was added to the solution, followed by 1 h incubation at 42°C. The temperature was then raised to 70°C for 10 min to heat inactivate Superscript II.

#### Second strand synthesis

49 μL of the second strand mix, containing 33.5 μL water, 12 μL 5x second strand buffer [100 mM Tris-HCl (pH 6.9), 23 mM MgCl2, 450 mM KCl, 0.75 mM β-NAD, 50 mM (NH4)2 SO4, Invitrogen, Cat. # 10812-014], 1.2 μL 10 mM dNTPs, 0.4 μL *E. coli* ligase (Invitrogen, Cat. # 18052-019), 1.5 μL DNA polymerase I (Invitrogen, Cat. # 18010-025), and 0.4 μL RNase H (Invitrogen, Cat. # 18021-071), was added to the product from the previous step. The mixture was incubated at 16°C for 2 h. cDNA was purified with 1x AMPure XP DNA beads (Beckman Coulter, Cat. # A63881) and eluted in 24μL nuclease-free water that was subsequently concentrated to 6.4 μL.

#### *In vitro* transcription

The concentrated solution was mixed with 9.6 μL of Ambion *in vitro* transcription mix (1.6 μL of each ribonucleotide, 1.6 μL 10x T7 reaction buffer, 1.6 μL T7 enzyme mix, MEGAscript T7 Transcription Kit, Thermo Fisher Scientific, Cat. # AMB13345) and incubated at 37°C for 13 h. Next, the aRNA was treated with 6 μL EXO-SAP (ExoSAP-IT™ PCR Product Cleanup Reagent, Thermo Fisher Scientific, Cat. # 78200.200.UL) at 37°C for 15 min followed by fragmentation with 5.5 μL fragmentation buffer (200 mM Tris-acetate (pH 8.1), 500 mM KOAc, 150 mM MgOAc) at 94°C for 3 min. The reaction was then quenched with 2.75 μL stop buffer (0.5 M EDTA) on ice. The fragmented aRNA was size selected with 0.8x AMPure RNA beads (RNAClean XP Kit, Beckman Coulter, Cat. # A63987) and eluted in 15 μL nuclease-free water. Thereafter, Illumina libraries were prepared as described previously [20].

### EMBR-seq with TEX digestion

To test the Terminator^™^ 5’-phosphate-dependent exonuclease (Lucigen, Cat. # TER5120), 100 ng of total RNA in 2 μL was combined with 18 μL TEX mix, comprised of 14.5 μL nuclease free water, 2 μL Terminator 10x buffer A, 0.5 μL RNAseOUT, and 1 μL TEX. The solution was incubated at 30°C for 1 h and quenched with 1 μL of 100 mM EDTA. The product was purified with 1x AMPure RNA beads and eluted in 10 μL nuclease-free water and concentrated to 2 μL. This TEX digested total RNA was then used as starting RNA in the EMBR-seq protocol described above.

### Bioinformatic analysis

Paired-end sequencing of the EMBR-seq libraries was performed on an Illumina NextSeq 500. All sequencing data has been deposited to Gene Expression Omnibus under the accession number GSE149666. In the sequencing libraries, the left mate contains information about the sample barcode (see *Primers*). The right mate is mapped to the bacterial transcriptome. Prior to mapping, only reads containing valid sample barcodes were retained. Subsequently, the reads were mapped to the reference transcriptome (*E. coli* K-12 substr. MG1655 cds ASM584v2) using Burrows-Wheeler Aligner (BWA) with default parameters.

### Primers

Reverse transcription primers are shown below with the 6-nucleotide sample barcodes underlined [20]:

GCCGGTAATACGACTCACTATAGGGAGTTCTACAGTCCGACGATCNN NNN N<u>(NNNNNN)</u>TTT TTTTTTTTTTTTTTTTTTTTTV

The following five barcodes were used in this study:

<u>AGACTC</u>
<u>AGCTTC</u>
<u>CATGAG</u>
<u>CAGATC</u>
<u>TCACAG</u>

Blocking primers:

5S 5’-ATGCCTGGCAGTTCCCTACTCTCGCATGGG-3’
16S 5’-TAAGGAGGTGATCCAACCGCAGGTTCCCCT-3’
23S 5’-AAGGTTAAGCCTCACGGTTCATTAGTACCG-3’

In the case of the 3’ phosphorylated primers, all blocking primers have a 3’ phosphorylation modification.

## Availability of data and materials

The datasets generated and analyzed in the current study have been deposited to GEO under the accession number GSE149666.

## Authors’ contributions

C.W. and S.S.D. conceived the method, C.W., K.A.H. and S.S.D designed the experiments. K.A.H., C.W. and E.L. participated in data collection. K.A.H. and C.W. analyzed the data. C.W., K.A.H., M.A.O. and S.S.D wrote the manuscript. S.S.D guided experimental design and data analysis. All authors have read and approved the final version of the manuscript.

## Acknowledgements

We would like to thank members of the Dey and O’Malley groups for helpful discussions. The authors thank Dr. Jennifer Smith for assistance with sequencing Illumina libraries. Sequencing was performed at the Biological Nanostructures Laboratory within the California NanoSystems Institute (CNSI), supported by the University of California, Santa Barbara (UCSB) and the University of California, Office of the President (UCOP). We further acknowledge the Center of Scientific Computing at UCSB for computational facilities that are funded by NSF MRSEC (DMR-1720256) and NSF CNS-1725797.

