## Supplementary Information for "Efficient and cost-effective bacterial mRNA sequencing from low input samples through ribosomal RNA depletion"

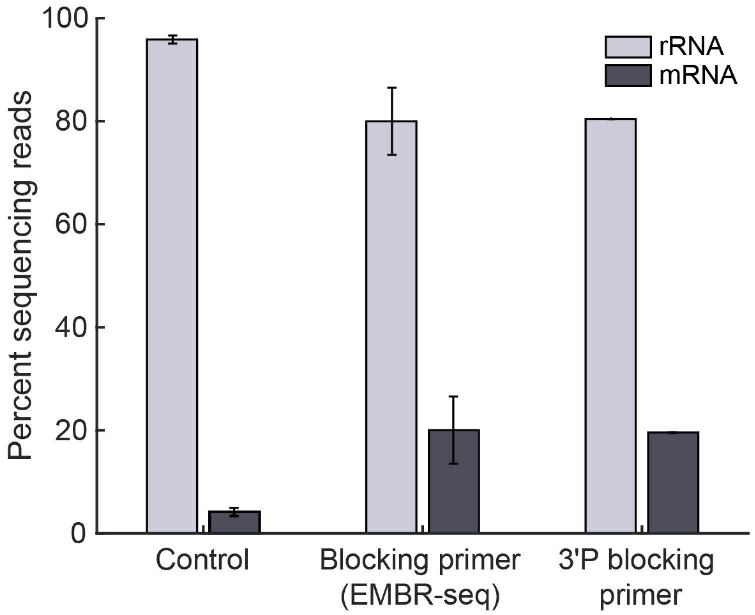


**Figure S1: Combining Terminator™ 5'-phosphate-dependent exonuclease (TEX) digestion with EMBR-seq does not improve rRNA depletion.** Performing TEX digestion prior to EMBR-seq results in less efficient rRNA depletion and mRNA enrichment compared to experiments without TEX (Fig. 2a) ($n \geq 2$, except in TEX + 3'P blocking primer where $n = 1$). These experiments were performed starting with 100 ng total RNA from *E. coli.* Error bars represent standard deviations.

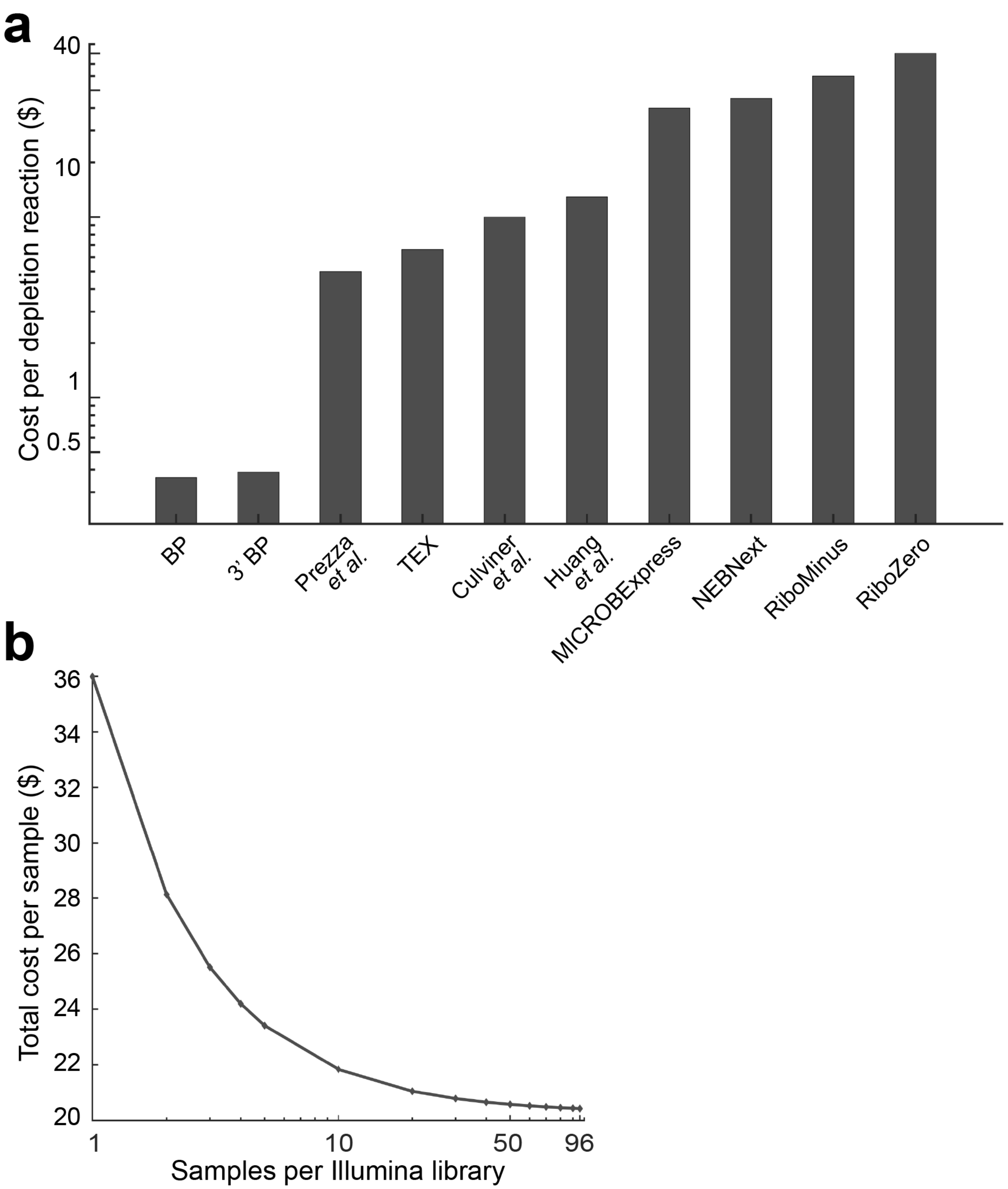


**Figure S2: Cost associated with performing EMBR-seq.** (**a**) The cost of performing rRNA depletion in EMBR-seq is ~$0.40 per reaction. The cost per reaction in EMBR-seq is an order of magnitude lower than other published rRNA depletion methods and commercial kits (Supplementary Table 1). (**b**) The plot shows the total cost for the complete EMBR-seq protocol per sample (starting from total bacterial RNA extraction to Illumina library preparation) as a function of the number of samples multiplexed (using the sample barcodes in the RT primer) per Illumina library. Starting from 1 sample per Illumina library to 96 samples per Illumina library, the total cost drops from $36 to $20 per sample.

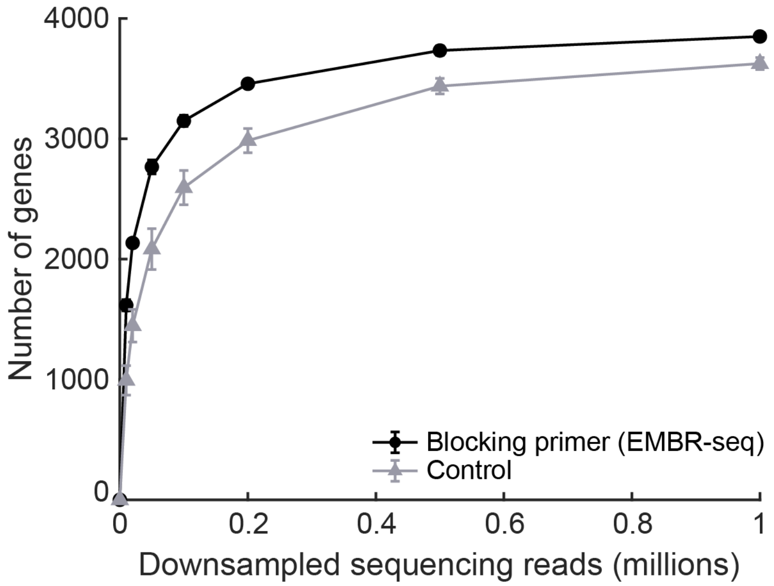


**Figure S3: Higher number of genes detected in EMBR-seq is not dependent on the sequencing depth.** To ensure that the number of genes detected in EMBR-seq samples compared to the control samples is not an artifact of sequencing depth, we downsampled the mapped sequencing reads to show that EMBR-seq detects more genes at different levels of downsampling. The figure also shows that the number of genes detected does not increase substantially beyond ~0.5 million mapped reads, suggesting that our sequencing libraries have been sequenced at sufficient depth ($n = 3$). Error bars represent standard deviations. For the EMBR-seq group, error bars are of the same scale as the size of the data points.

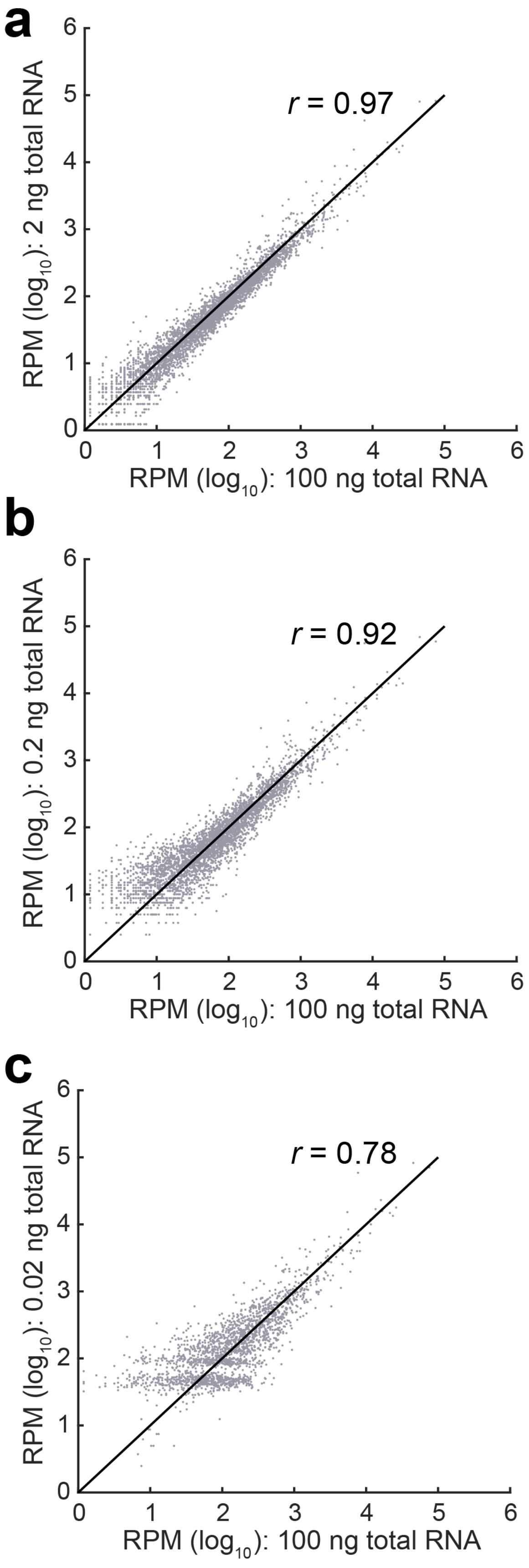
a
r = 0.97
RPM (log10): 2 ng total RNA
RPM (log10): 100 ng total RNA
b
r = 0.92
RPM (log10): 0.2 ng total RNA
RPM (log10): 100 ng total RNA
c
r = 0.78
RPM (log10): 0.02 ng total RNA
RPM (log10): 100 ng total RNA

**Figure S4: Gene transcript count correlation between different input total RNA amounts in EMBR-seq.** (**a**-**c**) Panels show gene transcript count Pearson correlations between 100 ng starting total RNA and lower input total RNA in EMBR-seq. As expected, the Pearson correlation drops when starting with lower amounts of total RNA. These experiments were performed with total RNA from *E. coli.*
